# Collaborations and deceptions in strategic interactions revealed by hyperscanning fMRI

**DOI:** 10.1101/2021.07.11.451985

**Authors:** Siao-Shan Shen, Jen-Tang Cheng, Yi-Ren Hsu, Der-Yow Chen, Ming-Hung Weng, Chun-Chia Kung

## Abstract

Despite its ubiquity, deceiving as a social phenomenon is scarcely addressed with fMRI, partly due to the spontaneity and individual differences in cheating, and the contextual variability that fosters lying. In this hyperscanning fMRI study, the participant pairs (n=33) from Taipei and Tainan joined an opening-treasure-chest (OTC) game, where the dyads took alternative turns as senders (to inform) and receivers (to decide) for guessing the right chest. The cooperation condition was achieved by, upon successful guessing, splitting the $200NTD trial reward, thereby promoting mutual trust. The competition condition, in contrast, was done by, also upon winning, the latter receivers taking all the $150NTD reward, thereby encouraging strategic interactions. One key fMRI finding was the negative correlations between the connectivity of the right temporo-parietal junction (rTPJ), known as the theory-of-mind function, and amygdala, parahippocampal gyrus, and rostral anterior cingulate (rACC), to senders’ behavioral lying rates. Furthermore, the Multi-Voxel Pattern Analysis (MVPA) over multiple searchlight-identified Region-Of-Interests (ROIs), in classifying either the “truthful vs. lying in $150” or the “truthful in $200 vs. truthful in $150” conditions achieved 61% and 84.5% accuracy, respectively, reflecting the idiosyncratic brain networks involved in distinguishing the social trust vs. deceptions in the dyadic interactions.

## Introduction

In modern societies, people interact with a variety of reasons or motivations. Collaborations and deceptions are two common scenarios where each individual’s underlying motivations may vary, but the aims of obtaining personal gains or advantages on one’s own side remain consistent. Social neuroscience, the research discipline that targets the brain mechanisms mediating social interactions, has kept keen interest in these two central social phenomena (for reviews, see Olsson, Knapska, & Lindstrom, 2020; Redcay & Schilbach, 2019). Also partially influenced by early optimism of treating fMRI ‘the next candidate of lie detector’(Farah, Hutchinson, Phelps, & Wagner, 2014), the neuroimaging research community have tried several paradigms to study the neural mechanisms underlying truth telling and, especially, deception.

To date, the fMRI literature on truth-telling and deception can be broadly categorized as ‘social vs. non-social interaction’. As shown in Fig. 1 for better illustration, within non-social interactions there are ‘instructed lying’ (instructing subjects to tell lies in specific conditions/blocks) and ‘spontaneous lying’ (designing incentives so that participants may or may not lie in a natural manner), containing several tasks for each. For example, in instructed lying, when later facing previously presented vs. unpresented words (or objects) in the recognition phase, participants were instructed to lie (e.g., saying ‘new’ to old/familiar items) or to tell the truth (Abe et al., 2008); or similar responses (e.g., instructed to judge the previously presented faces as ‘unfamiliar’) ‘in the guilty knowledge test (Gamer, Bauermann, Stoeter, & Vossel, 2007). However, people lie in daily lives as voluntary or spontaneous, so the findings from instructed lies may be ‘lacking ecological validity’ (e.g., pp. 1102, in Yin & Weber, 2019). In spontaneous lying paradigms, in contrast, experimenters designed novel or creative contexts (see Fig. 1 for the listed games in the category) where incentives, such as attractive monetary rewards (Yin, Reuter, & Weber, 2016; Yin & Weber, 2019), or appearing ‘uncaught-ness’ (Speer, Smidts, & Boksem, 2020), to induce participants to spontaneously lie.

**Figure 1:**
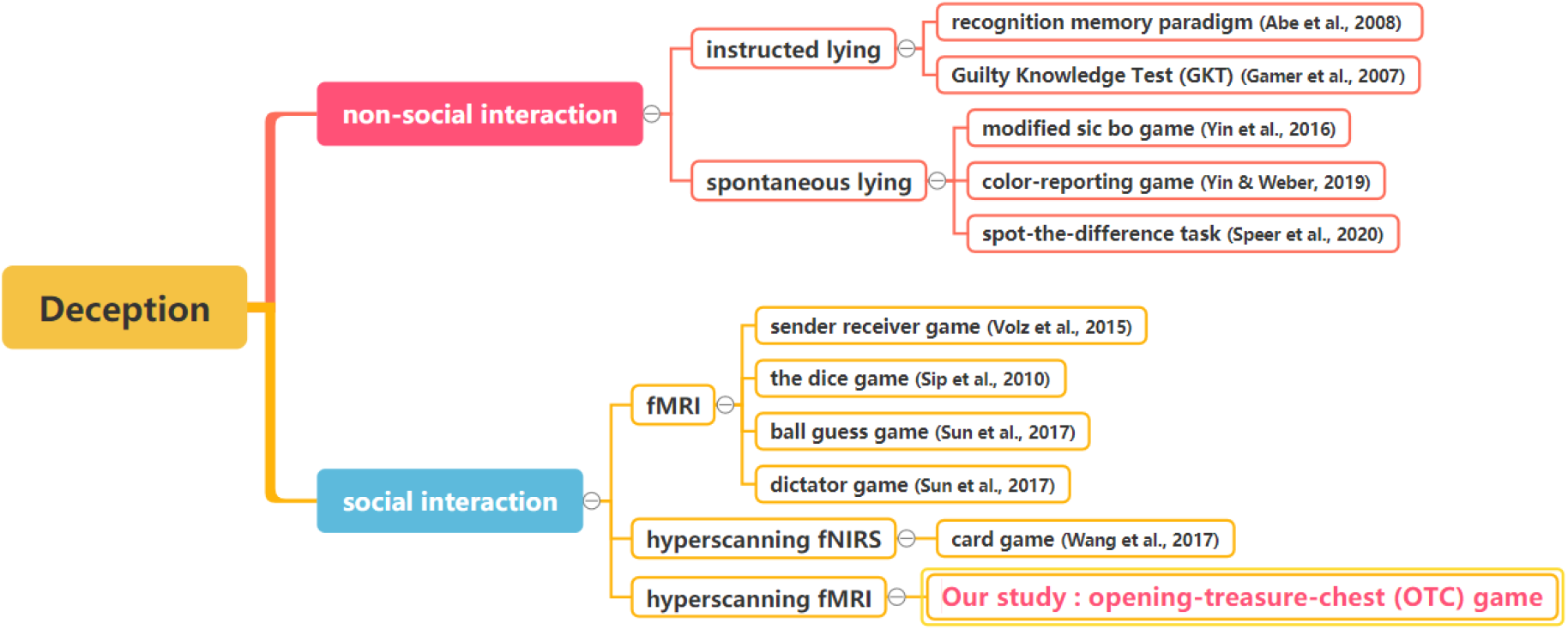
Selected hyperscanning neuroimaging papers that involve deception. Highlighted are the category (non-social vs. social), imaging facility: fMRI and fNIRS; and most importantly, the tasks that facilitated or created the deception context. What distinguished non-social (red) and social (blue patch) paradigms is whether the task involves human to agent (or human) interactions.

The lower half of Fig. 1, shown in blue patch, lists several neuroimaging studies (mostly fMRI and fNIRS) bearing ‘social’ deception. For example, the Bluff Card Game (Wang, Wang, Zhou, & Yu, 2020; Zhang, Liu, Pelowski, & Yu, 2017) mimics the real life poker game, where players may occasionally bet with more tokens despite bad cards. In another study, Volz, Vogeley, Tittgemeyer, von Cramon, and Sutter (2015), subjects were told to interact with the other subject (but in reality a computer program, unbeknown to the participants) in a sender-receiver paradigm, where one instructed the imaginary other for the competitive reward, thereby motivating the social interactions of truth telling and misleading behaviors. However, as Fig. 1 also shows, that most of the cited fMRI studies on deception, though social in nature (e.g., person-agent/program interactions), only involved a single MRI machine; whereas the real hyperscanning studies, involving two or more machines/facilities simultaneously by definition (Czeszumski et al., 2020; Montague et al., 2002), on deception all adopted functional near-infrared spectroscopy (fNIRS), measuring surface cerebral cortices up to 1.5cm depth (Quaresima & Ferrari, 2016). To our knowledge, the fMRI hyperscanning study on deception has yet to be reported.

The above-mentioned neuroimaging studies have gathered the suggested brain networks involved in deception, which can be broadly categorized as the three functional subsystems: (a) including the dorsolateral prefrontal cortex (dlPFC), the ventrolateral prefrontal cortex (vlPFC), the inferior parietal lobule (IPL), the inferior frontal gyrus (IFG), the superior frontal gyrus (SFG), the anterior cingulate cortex (ACC), insula, and other related networks. Most of these regions are associated with executive control (Cui et al., 2014; Ito et al., 2012; Sun et al., 2017), inhibition control (Vartanian et al., 2013; Zhu et al., 2014), conflict monitoring (Yin et al., 2016), or emotion regulation (Suzuki, Misaki, Krueger, & Bodurka, 2015; Yin, Hu, Dynowski, Li, & Weber, 2017); (b) the Theory of Mind (ToM) (Premack & Woodruff, 2010) network also plays an important role in social deception, involving right temporal-parietal junction (rTPJ), the medial prefrontal cortex (mPFC), and the posterior cingulate cortex (PCC) or precuneus (Schurz, Radua, Aichhorn, Richlan, & Perner, 2014). Among them, the rTPJ involves taking another person’s perspective and moral reasoning (Saxe & Kanwisher, 2003; Saxe & Wexler, 2005), therefore commonly adopted as the seed base for advanced analyses; (c) the reward-related processing: such as the nucleus accumbens (NAcc), caudate, putamen, and the ventromedial prefrontal cortex (vmPFC), commonly associated with monetary reward and value coding (Abe & Greene, 2014; Pornpattananangkul, Zhen, & Yu, 2018; Yin & Weber, 2019). Despite the accumulated findings on deception, the findings are not consistent across studies, probably due to the experimental and individual differences. To illustrate this point, consider what was routinely mentioned in related review articles, the famous opposing-viewpoint theories: Grace vs. Will hypotheses (Greene & Paxton, 2010). The Grace hypothesis proposes that people are men of good nature. They need to override their default honest behavior, so the cognitive control networks are required to inhibit impulses for the purpose of profit. The study of Sun et al. (2017) revealed that the cognitive and inhibition control regions showed greater activation for deception. On the other hand, the Will hypothesis suggested that people’s nature is evil at birth, and their honest decisions are the result of resisting temptation. Once the activation of cognitive control regions such as the dlPFC decreases, the lower honesty concerns are the consequences of economic games (Zhu et al., 2014). These two competing hypotheses can therefore be the result of individual differences in the moral identity (Speer et al., 2020).

Besides paradigmatic and contextual differences, another layer of complexity that accompanied the relative scarcity of hyperscanning neuroimaging studies, is the relative limited methodologies adopted: most cited fMRI studies deployed univariate (e.g., general linear model or GLM) or region-of-analysis methods (e.g., Abe & Greene, 2014; Ito et al., 2012; Lisofsky, Kazzer, Heekeren, & Prehn, 2014; Sun et al., 2017; Vartanian et al., 2013; Wu, Loke, Xu, & Lee, 2011; Yin et al., 2016), with few recent exceptions utilizing connectivity (Jiang et al., 2015; Speer et al., 2020) and multivariate approaches (Jin et al., 2009; Yin & Weber, 2019). In comparison, in the present study, GLM contrast, psychophysiological interactions (PPI, O’Reilly, Woolrich, Behrens, Smith, & Johansen-Berg, 2012), and multi-voxel pattern analysis (MVPA, Kriegeskorte, Goebel, & Bandettini, 2006) were adopted to identify patterns of task- or seed-driven activity throughout the brain, plus the subsequent principal component analysis (PCA) that reduced the high-dimensionality problems in MVPA analysis, expanding the perspectives of data examination and joint constraints of the result interpretations.

Given above, the current study aims to investigate the neural mechanisms of interpersonal collaborations and deceptions, with an Opening Treasure Chest (OTC) game under the fMRI hyperscanning setup. The OTC game is a strategic game based on the sender-receiver paradigm, consisting of two conditions: cooperation and competition. To realize deception, participants (e.g., the sender) in the current study’s competition condition could mislead opponents (e.g., the receiver) with true/false information about the location of the treasure, for reward maximization. The truth-telling strategy was operationally defined by senders’ suggestions of the location of the treasure chests with higher probability, whereas the lying strategy by senders’ suggesting the low probability side. The expected neural mechanism under deception may involve three cognitive systems, as Sip, Roepstorff, McGregor, and Frith (2008) identified: executive and inhibition control (e.g., coordinating multiple sources of information), mentalizing (e.g., perspective-taking; inferring mental states of others, etc.), and reward processing (reward anticipation and/or feedback update, etc.) networks. In addition, because connectivity-based studies have shown that r-TPJ, the important site of mentalizing others, connected more to self-related processing regions, such as medial prefrontal (mPFC) and precuneus (Bitsch, Berger, Nagels, Falkenberg, & Straube, 2018), during competition than during coordination conditions, in our similar psychophysiological interaction (PPI) analyses the same expectation could also be drawn. As far as MVPA results are concerned, one would predict the similar brain areas revealed by both GLM contrast and MVPA searchlight mappings on identical contrasts (Davis et al., 2014). Besides, other studies (Tsoi, Dungan, Waytz, & Young, 2016) have shown that one can paternal differences in brain areas not differentiable by GLM contrast, e.g., the truth-telling vs. lying strategies in competition conditions, could be separated by MVPA.

## Materials and methods

### Participants

Sixty-six (33 pairs) participants, between 20 and 30 years of age (M=23.4, SD=2.9), participated in the study. They were recruited from National Taiwan University (NTU), Taipei, and National Cheng Kung University (NCKU), Tainan. All participants were native Taiwanese speakers, who had normal or corrected-to-normal vision, and reported no history of psychiatric or neurological disorders. All methods and procedures were performed following the relevant guidelines and regulations approved by the NCKU Governance Framework for Human Research Ethics https://rec.chass.ncku.edu.tw/en, with the case number 106-254. After the fMRI experiment, each participant received 600 $NTD, plus the bonus calculated by the sum of won money divided by 112 (total trial number), around ~300 $NTD.

### Experimental task

In the present study, the opening-treasure-chest (OTC) game was carried out in a hyperscanning fMRI context. Participants were told that they would interact with a real partner from another school to play the OTC game together. The game has both the cooperation and the competition condition (see Fig. 2). In each trial, there were two treasure chests presented on the screen, with only one chest containing money (and the other empty). The subjects were told that their goal was to guess the right treasure chest and win the reward. When being the sender, only he/she was informed about the probabilities of money being in the left and right chest, and the sender’s role was to suggest which chest the receiver should choose. After receiving the suggestion from the sender, the receiver decided which chest to open. In the cooperation condition, the treasure chest chosen by the receiver was the final choice of both the sender and the receiver, the dyad then split the $200 reward if they successfully guessed the correct treasure chest; whereas in the competition condition, after the receiver decided which treasure chest to open, the sender was automatically assigned to open the other treasure chest, and the person who chose the correct chest took all the $150-reward. In the collaboration condition, the program was set to 75% success rate (meaning that one out of every 4 trials will show up on the low probability side), so that each player’s expected utility for each trial was 75 $NTD. In the competition condition, the success rate was set to be 50%-50% of the final treasure chest, therefore the final expected utility for either participant was also 150 x 50% = 75 $NTD.

**Figure 2.**
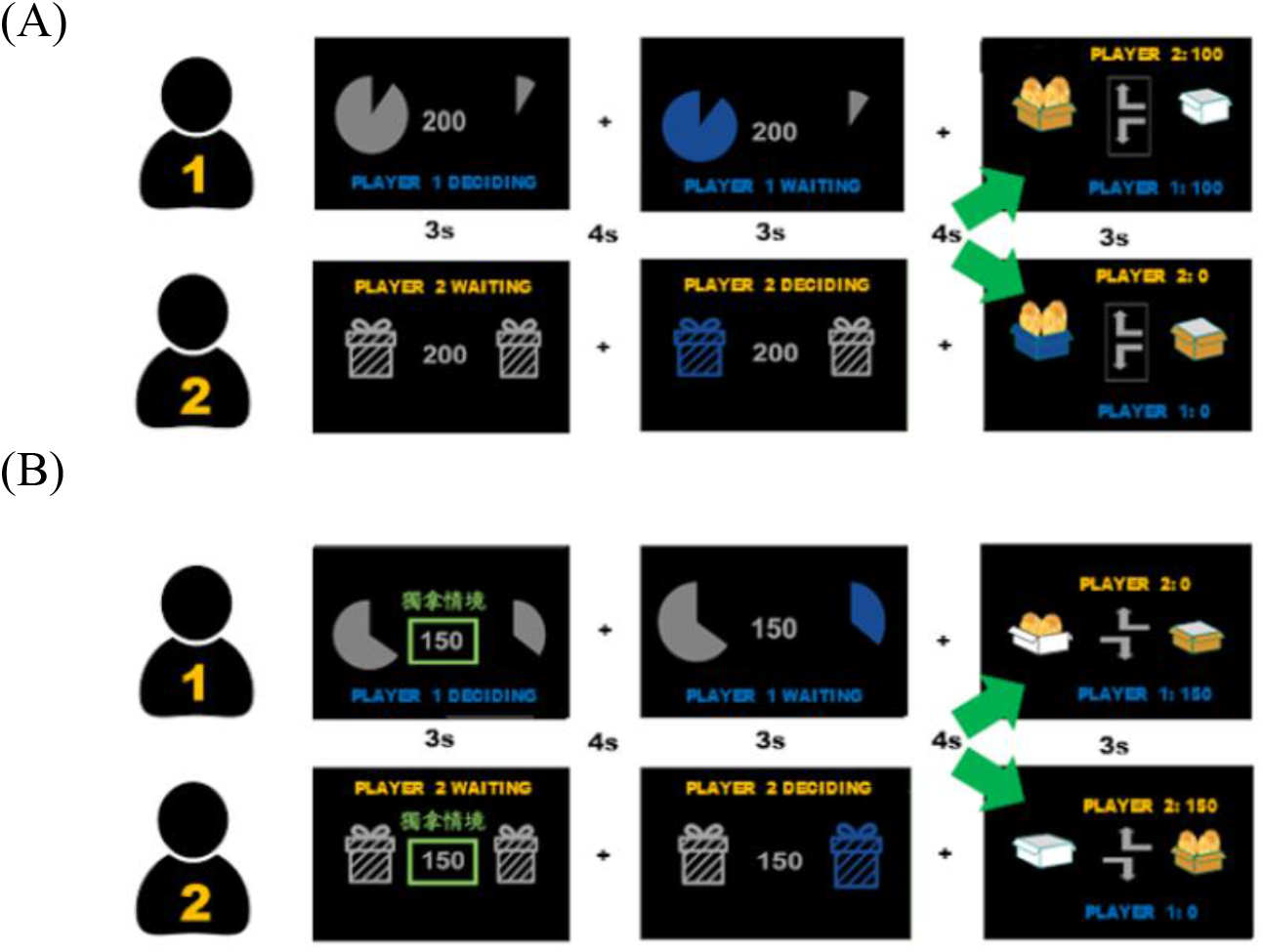
Illustrations of the hyperscanning fMRI experiment of the strategic (e.g., Opening Treasure Chest, or OTC) game, with two conditions. (A) In the cooperation case ($200), the dyad split the reward if they succeeded in guessing the correct treasure chest (e.g., the receiver followed the sender’s left/right signal). (B) In the competition case ($150), either the sender or the receiver took all the money (e.g., the sender suggested the right chest, and the receiver chose the left one and won). Trials were in the fixed order of ‘$200-sender, $200-receiver, $150-sender, $150-receiver’ units, repeated 7 times per run.

After some practices and initial setup and hyperscanning testing, both participants completed 4 functional runs of 28 trials in each, during which the 4-trial segments: $200-sender, $200-receiver, $150-sender, and $150-receiver, would repeat 7 times. Which site to be the first sender role was determined randomly. The subjects were allowed to have a short break between each run, and the game resumed when both players were ready to continue. Each trial took approximately 17 seconds to complete: three seconds for the sender’s decision, four seconds of fixation, three seconds for the receiver’s decision, four seconds of fixation, and three seconds of feedback respectively. The inter-trial interval was approximately 3 to 9 seconds. Each pair of participants took approximately 70 minutes to finish, given that we typically ran 3 pairs in the 4-hr (Monday 9am-1pm) slot.

### fMRI acquisition and preprocessing

fMRI data were acquired with a 3 T General Electric 750 MRI scanner (GE Medical Systems, Waukesha, WI) located in the B3F of the NCKU MRI center (http://fmri.ncku.edu.tw), equipped with a standard 8-channel head coil. Whole-brain functional scans were acquired with a T2* EPI (TR = 2s, TE = 33ms, flip angle = 90 deg, 40 axial slices, voxel size = 3.5 × 3.5 × 3 mm^3^). High-resolution T1-weighted structural scans were acquired using a 3D fast spoiled grass (FSPGR) sequence (TR = 7.65 ms, TE = 2.93 ms, inversion time = 450 ms, FA= 12 degree, 166 sagittal slices, voxel size = 0.875 × 0.875 × 1 mm^3^). Another scanner is the 3-Tesla PRISMA MRI (Siemens, Erlangen, Germany) located at the National Taiwan University (http://mrimeg.psy.ntu.edu.tw/doku.php), equipped with a 20-channel phase array coil. Whole-brain functional scans were acquired with a T2* EPI (TR = 2s, TE = 24 ms, flip angle = 87 degree, 36 axial slices, voxel size = 3 × 3 × 3 mm^3^). High-resolution structural scans were acquired using a T1-weighted (MP-RAGE) sequence (TR = 2s, TE = 2.3 ms, inversion time = 900ms, FA= 8 deg, 192 sagittal slices, voxel size = 0.938 × 0.938 × 0.94 mm^3^). The fMRI data were preprocessed and analyzed using BrainVoyagerQX v. 2.6 (Brain Innovation, Maastricht, The Netherlands), NeuroElf v1.1 (https://neuroelf.net) under Matlab 2019a (MathWorks, Natick, MA). After slice timing correction, functional images were corrected for head movements using the six-parameter rigid transformations, aligning all functional volumes to the first volume of the first run. High-pass temporal filtering (with the default BVQX option of GLM-Fourier basis set at 2 cycles per deg, but no spatial smoothing) was applied. The resulting functional data were co-registered to the anatomical scan via initial alignment (IA), final alignment (FA), and then both the functional (*.fmr) and anatomical (*.vmr) files were transformed into the Talairach space.

### Behavioral data analysis

Behavioral data, primarily the reaction times during decision phase, were analyzed with Excel (2013) and JASP 0.11.1.0 (https://jasp-stats.org/). One-way ANOVAs and paired t-tests were applied to compare the effect of subjects’ decisions in cooperation vs. competition conditions, and during interaction phases (sender vs. receiver decision times).

### fMRI data analysis

Like the majority of task-based fMRI papers, the first step of the analysis is usually the general linear model (GLM), estimating each participant’ brain activations and comparing the contrasts of interests under different stages of decision-making. Six regressors of interest were included in the GLM: (a) senders’ decision phases in the cooperation condition; (b) senders’ decision phases in the competition condition; (c) receivers’ decision phases in cooperation; (d) receivers’ decision phases in competition; (e) feedback stages in cooperation; and finally, (f) feedback stages in the competition condition. For the second level (random-effect) analysis, three different contrasts were explored (and offered in results): competition versus cooperation in the sender decisions, the comparison of truth-telling strategy (e.g., the sender suggests the receiver the chest with higher probability) under the corporation vs. under the competition conditions, and the truth-telling vs. lying strategies (e.g., the sender suggests the receiver the chest with lower probability) under the competition condition. Because the goal of the present study is on social deception, senders’ decision under both conditions will be the primary focus.

In addition to the GLM contrasts, the psychophysiological interactions (PPI) analyses were applied to examine differences of functional connectivity between seed region(s) of interest and target ROIs. The seed regions included the right TPJ, bilateral dorsolateral PreFrontal Cortex (DLPFC) and bilateral caudate. These seeds were downloaded from the online meta-database Neurosynth (https://neurosynth.org/) as .nii masks. In addition, to investigate the individual differences in the ratio of truth-telling to lying, and such relationship with their functional connectivity, participants’ percentage of lying strategies was correlated with their connectivity differences to reveal behavior-relevant brain networks.

Furthermore, to explore the cognitive states under the competition or cooperation conditions, where the participants’ strategies varied, a multi-voxel pattern analysis (MVPA) (Kriegeskorte et al., 2006) whole-brain searchlight mapping, combining searchmight toolbox (http://www.franciscopereira.org/searchmight/) and the Princeton MVPA toolbox (http://code.google.com/p/princeton-mvpatoolbox/), to classify the patterns of neural activity under conditions of interest or under senders’ decisions. In the searchlight mapping analyses, a Gaussian Naïve Bayesian (GNB) classifier, plus the leave-one-TrialPair-out cross-validation scheme, were to calculate each participant’s classification accuracy under conditions of interest. Since each sender yielded different numbers of truth-telling vs. lying decisions, the trial numbers entered the MVPA needed to be down sampled to balance the training and testing data sets. Further multi-ROI MVPAs, adding up to 10 ROIs derived from the searchlight-mapping clusters, were calculated (Chou et al., 2014) to assess the increase of classification performances, under the truth-telling versus lying strategy in competition and under the truth-telling strategies in competition versus cooperation conditions. Finally, to reduce the influence of hyper-dimensional collinearities the voxels/features shared in the MVPA analysis, dimensionality reduction techniques, such as the principal component analysis (PCA) (Kim et al., 2017) was used to convert high dimensional neural representations into principal components, to evaluate whether there were improved performances.

## Results

### Behavioral Results

Across all 66 participants, truth-telling strategies in senders’ cooperation conditions were averagely adopted 98.2 ± 2.8%, ranged from 88% to 100% of the time; whereas in the competition condition, this percentage dropped to 54.5±17%, ranged from 18% to 100% and the lying strategies 45.5 ± 17%, ranged from 82.1% to 0% (Fig. 3A). The receivers’ complementary ‘following’ rate (e.g., when the senders suggested left, the receivers responded accordingly) were: 98.7% in the cooperative condition (SD=2.4%, from 89.3 to 100%), and in the competition condition, the following rate was 57.9 ± 14.5%, from 21.4 to 92.9%, and ‘unfollowing’ rate 42.1±14.5%, from 78.6 to 7.1% (Fig. 3B). Overall, our task manipulations achieved the desirable outcome: each pair cooperated exclusively in the cooperation condition; and mixed their truth-telling and lying strategies, and the correspondingly following vs. unfollowing behaviors, evenly in the competition condition.

**Figure 3.**
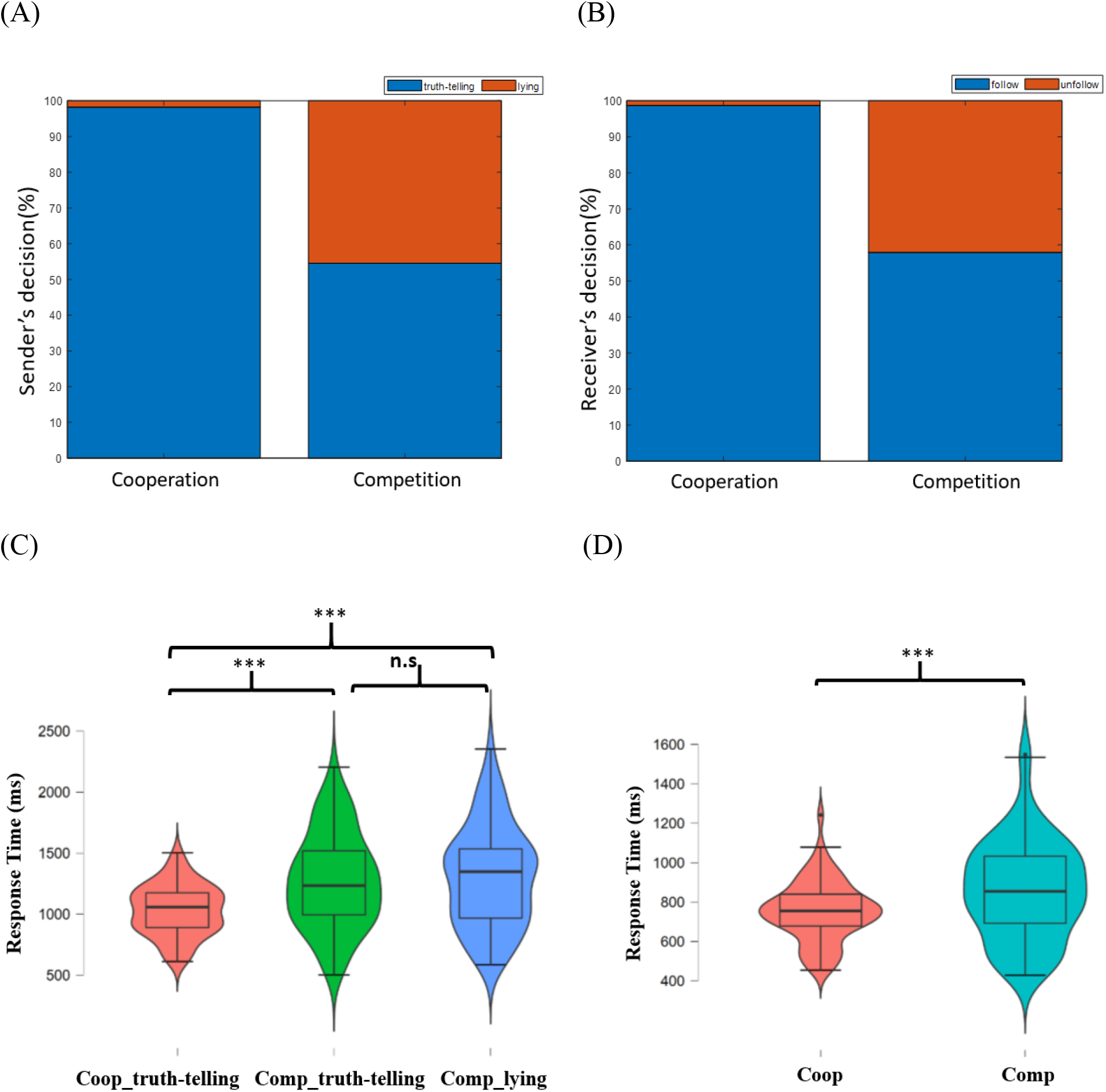
(A&B) The mean proportions (N=66) in “truth-telling/blue-color (defined as “seeing left and pointing left) vs. lying (“seeing left and pointing right)” behaviors for senders (A), and “follow/blue-color vs. unfollow” for receivers (B). (C&D): the mean RT results of 64 participants (2 excluded) in 3 conditions. (C) The mean response times in the competition and the cooperation conditions during senders’ decision phase; (D) The mean response times in the competition and the cooperation conditions during receivers’ decision phase. (*** p < .001; n.s.: not significant).

Among the 64 participants included in the statistical analyses (two subjects were excluded due to their anonymously truth-telling strategies in the sender condition), the mean response time (RT) of the three following scenarios: (1) & (2) are truth-telling and lying in the competition condition, respectively; and (3) truth-telling in the cooperation condition. Lying choices by the senders were defined by their suggesting the receivers to choose the chest of lower probability, whereas senders’ truth-telling choices suggested the chest with higher probability. In the cooperation condition, in contrast, truth-telling decisions reflected the majority of senders’ suggesting the receivers to choose the chest with high probability. Across the competition condition, the mean response time (RT) was 1,217 ms (SD = 49 ms) for truth-telling, and 1306 ms (SD = 51 ms) for lying; and the RT for truth-telling trials was 1031 ms (SD = 26 ms) in the cooperation condition. The one-way ANOVA on these RTs showed that they were significantly different from the all-equal null hypothesis (F _(2,189)_ = 11.82) (see Fig. 3C). The post hoc analysis on the truth-telling trial RTs of the cooperation condition were significantly less than those of both the truth-telling (t_(63)_ = 3.907, p < .001) and lying (t_(63)_ = 4.46, p < .001) trials in the competition condition, but no significant difference between lying and truth-telling RTs of the competition condition (t_(63)_ = 0.552, p = .845). The most plausible explanation, we surmise, is that the subjects in the competition condition, gradually realized that being honest or not did not guarantee higher winning probability, ended up spending equal amount of extra time in either truth-telling or lying, both longer than the RTs of plain mutual trusting in the collaboration conditions.

In the cooperation condition, the mean RT for the receiver to decide whether to follow the opponent’s suggestion was 754 ms (SD = 19 ms), while in the competition condition the mean RT was 859 ms (SD = 31 ms). The Student’s paired t-test results showed that the mean RT under the competition condition was longer than that under the cooperation condition (t _(63)_ = −4.914, p < .001) (Fig. 3D). The longer time receivers took in the competition conditions probably reflected their extra thought processes involved, such as whether to trust the senders or not, etc.

### fMRI Results

#### General Linear Model (GLM)

When comparing the sender’s brain activities during the decision phase under both the competition and the cooperation conditions, multiple areas showed higher brain activities in the former (e.g., competition): including the bilateral dlPFC, mPFC, IPL, IFG, insula, precuneus, caudate, rTPJ, etc. (see Fig. 4A, and Table 1 for comprehensive list).

**Figure 4.**
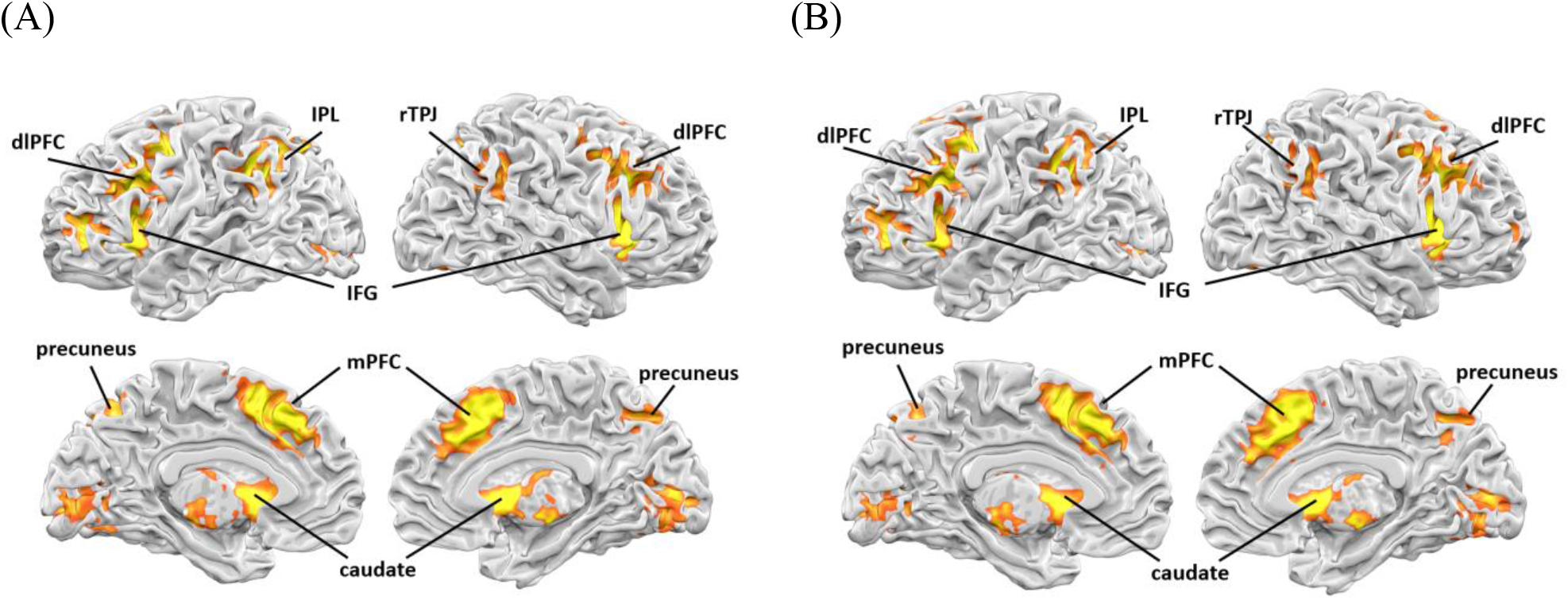
(A) GLM results of the contrast in the competition versus cooperation interactions during senders’ decision phase [Voxel-level threshold p < .005 uncorrected, (cluster threshold > 41 voxels, FWE p=.05 corrected thresholded), via Neuroelf’s Alphasim]; (B) GLM results of the contrast when the sender used truth-telling strategies in the competition and the cooperation [Voxel-level threshold p < .005 uncorrected, (cluster threshold > 33 voxels, FWE p=.05 corrected thresholded), via Neuroelf’s Alphasim] (also see Table 1&2). Note the high similarities among the activated regions/networks in both contrasts.

**Table 1.** The number of ROIs (with voxel numbers), their anatomical labels, xyz coordinates (Talairach space), and the associated t-values of over-thresholded clusters. Brain activation in the contrast of competition versus cooperation during the sender’s decision phase [Voxel-level threshold p < .005 uncorrected, (cluster threshold > 41 voxels, FWE p=.05 corrected thresholded), via Neuroelf’s Alphasim].

| Cluster | Anatomical region | x | y | z | t-value |
| --- | --- | --- | --- | --- | --- |
| Cluster1(Voxel : 637) | L Inferior Frontal Gyrus | -45 | 13 | 14 | 7.52 |
| Cluster2(Voxel : 211) | R Thalamus | 9 | 0 | 6 | 7.4 |
| Cluster4(Voxel : 708) | R Insula | 33 | 17 | 8 | 7.36 |
| Cluster5(Voxel : 739) | R Cingulate Gyrus | 3 | 19 | 37 | 7 |
| Cluster6(Voxel : 515) | L Middle Frontal Gyrus | -38 | 0 | 48 | 6.38 |
| Cluster7(Voxel : 497) | L Inferior Parietal Lobule | -41 | -46 | 44 | 6.2 |
| Cluster8(Voxel : 152) | L Inferior Frontal Gyrus | -50 | 39 | -1 | 5.23 |
| Cluster9(Voxel : 112) | L Precuneus | -13 | -68 | 41 | 4.68 |
| Cluster10(Voxel : 165) | R Middle Temporal Gyrus | 48 | -59 | 25 | 4.38 |
| Cluster11(Voxel : 77) | R Precuneus | 12 | -71 | 42 | 4.21 |

The neural activity between the truth-telling scenarios for cooperation and competition was compared, and the results showed that multiple brain areas had higher activities in the truth-telling trails of the competition condition than those of the cooperation condition. The brain areas included the bilateral caudate, insula, dlPFC, IPL, precuneus, mPFC, and the rTPJ (see Figure 4B, and Table 2).

**Table 2.**
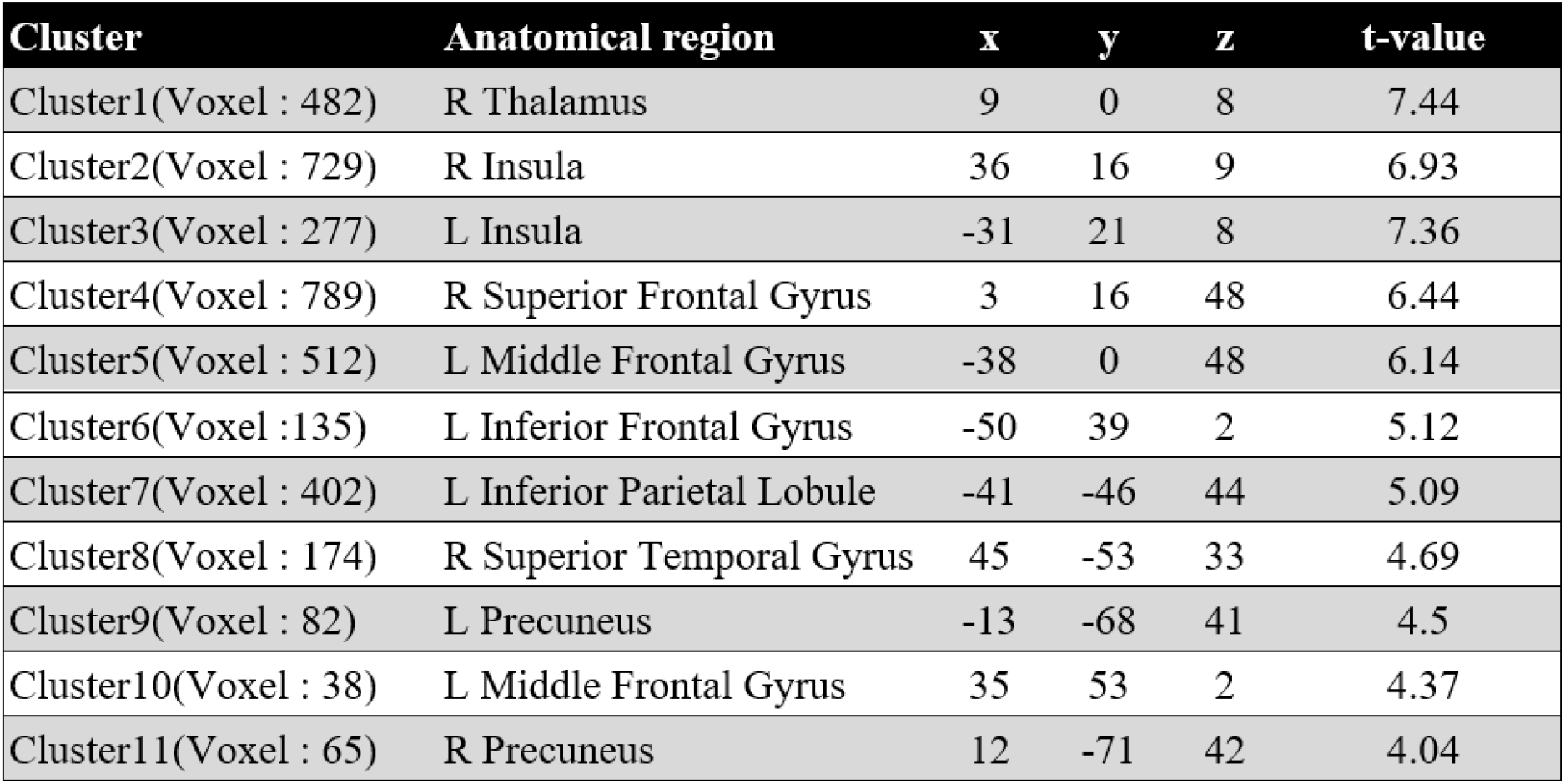
Brain activation in the contrast of truth-telling in competition versus cooperation [Voxel-level threshold p < .005 uncorrected, (cluster threshold > 33 voxels, FWE p=.05 corrected thresholded), via Neuroelf’s Alphasim].

A further analysis was conducted to compare the sender’s neural activity of truth-telling and lying strategies in the competition condition. However, no significant difference was found between these two conditions when participants were assigned as a sender. This result may be due to the fact that similar neural mechanisms were used in both strategies of truth-telling and lying in the competition condition. Together, the combined results of Fig. 4A (senders’ decisions between the competition and the cooperation conditions), 4B (senders’ truth-telling in cooperation vs. in competition condition), and the null results in the 3rd contrast (senders’ truth-telling vs. lying in the competition condition), mimicked the behavioral RT patterns, shown in Fig. 3C, very well, suggesting that these co-activated regions in both Fig. 4A and 4B contrasts, reflect common involvement of related networks (Sip et al., 2008), and that null results of ‘truth-telling vs. lying strategy in the competition condition’ (not shown) reflecting the similar undertakings in both lying and truth-telling, at least from the univariate analysis perspective.

### PsychoPhysiological Interaction (PPI)

The rTPJ, dlPFC and caudate were used as the seed regions to examine the target regions where their functional connectivity were significantly stronger in the competition vs. the cooperation conditions. As shown in Fig. 5, during the sender’s decision phase all three seeds were more connected to the bilateral precuneus, with the additional thalamus (Cunningham, Tomasi, & Volkow, 2017) more connected with both rTPJ and dlPFC seeds, and the left superior temporal gyrus (lSTG) solely with dlPFC, cuneus and left precentral gyrus with caudate, commonly in the “competition vs. cooperation” conditions (see Fig. 5 and Table 3).

**Figure 5.**
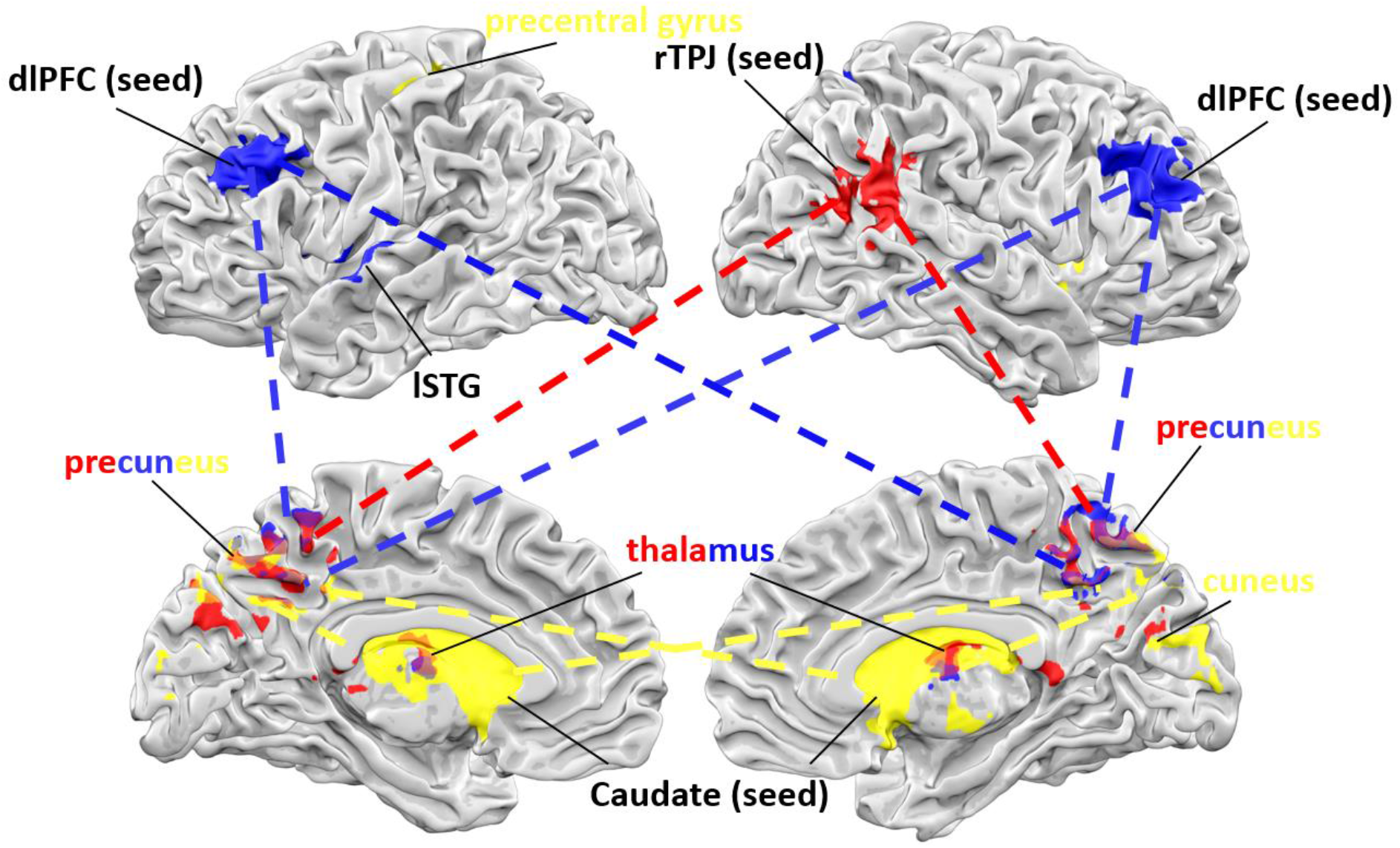
Results of the three seed-based PPI analyses comparing the competition vs. cooperation conditions. The three seed regions: rTPJ (red), dlPFC (blue) and caudate (yellow), during senders’ decision phase, commonly showed significantly higher connectivity (shown in dotted lines) to the bilateral precuneus. Likewise for the thalamus area that was commonly connected with both rTPJ and dlPFC seeds, and lSTG (shown in blue) singly with dlPFC, cuneus and l-precentral (shown in yellow) with caudate seed, during the same contrast and same decision phase. All these PPIs were done with the same voxel-level threshold p < .005 (uncorrected), cluster threshold > 22 voxels, FWE p=.05 (corrected), via Neuroelf’s Alphasim (also see Table 3).

**Table 3.** PPI analysis with three seed regions, including rTPJ, dlPFC and caudate, between the competition versus cooperation conditions, and during sender’s decision phase. [Voxel-level threshold p < .005 uncorrected, (cluster threshold > 22 voxels, FWE p=.05 corrected thresholded), via Neuroelf’s Alphasim].

| Cluster | Anatomical region | x | y | z | t-value |
| --- | --- | --- | --- | --- | --- |
| **rTPJ(seed)** |  |  |  |  |  |
| Cluster1(Voxel : 137) | L Precuneus | 0 | -47 | 36 | 5.9 |
| Cluster2(Voxel : 70) | R Thalamus | 0 | -10 | 14 | 4.84 |
| Cluster3(Voxel : 36) | R Thalamus | -6 | -34 | 7 | 4.77 |
| Cluster4(Voxel : 31) | L Precuneus | 0 | -67 | 23 | 4 |
| **dlPFC(seed)** |  |  |  |  |  |
| Cluster1(Voxel : 27) | L Thalamus | -3 | -9 | 12 | 4.83 |
| Cluster2(Voxel : 122) | L Precuneus | 0 | -56 | 32 | 4.62 |
| Cluster3(Voxel : 39) | L Superior Temporal Gyrus | -65 | -12 | 8 | 4.43 |
| **caudate(seed)** |  |  |  |  |  |
| Cluster1(Voxel : 32) | L Precuneus | 0 | -50 | 35 | 4.2 |
| Cluster2(Voxel : 25) | R Cuneus | 2 | -82 | 17 | 3.8 |
| Cluster3(Voxel : 60) | L Precuneus | 0 | -61 | 42 | 3.7 |
| Cluster4(Voxel : 26) | L Precentral Gyrus | -28 | -27 | 56 | 3.5 |

To investigate the underlying network differences when senders’ truth-telling strategies were in the competition vs. in the cooperation conditions, the rTPJ seed connectivity was further examined. The results, once again (very similar to Fig. 5 results in red colors), showed that higher functional connectivity was seen in the bilateral precuneus and thalamus when the senders chose to use the truth-telling strategy in competition than in cooperation (see Table 4). However, such PPI analyses with bilateral dlPFC and caudate seeds, under the same contrast and same thresholds, revealed none significant clusters whatsoever.

**Table 4.** PPI analysis with rTPJ seed regions. Functional connectivity in the contrast of competition versus cooperation when sender used truth-telling strategies [Voxel-level threshold p < .005 uncorrected, (cluster threshold > 20 voxels, FWE p=.05 corrected thresholded), via Neuroelf’s Alphasim].

| Cluster | Anatomical region | x | y | z | t-value |
| --- | --- | --- | --- | --- | --- |
| Cluster1(Voxel : 85) | R Precuneus | 3 | -48 | 36 | 5.29 |
| Cluster2(Voxel : 33) | L Thalamus | -6 | -34 | 7 | 4.84 |
| Cluster3(Voxel : 35) | L Thalamus | 0 | -10 | 12 | 4.39 |

Thirdly, the functional connectivity with all three seeds (rTPJ, bilateral-dlPFC and - caudate) showed consistently no significant clusters between senders’ truth-telling vs. lying strategies in the competition condition. With these similar findings between three types of GLM contrasts and the three types of corresponding PPI analysis between rTPJ and related networks, the implications strongly suggest that (a) in the cooperation condition, truth telling, which represented ~98% of the corporation trials, was synonymous to the whole cooperation condition; (b) the truth-telling and lying strategies in the competition conditions was, at least from the perspective of univariate analysis approach, also undifferentiated in terms of underlying neural substrates (given there was almost no way to separating these two conditions with the GLM contrasts, plus with the three different seed-based PPIs).

Fourthly, inspired by Hackel, Wills, and Van Bavel (2020), which employed the connectivity-based correlations with behavioral covariates, in the current study the similar connectivity-based correlations were also applied to predict participants’ strategies during competition conditions. Once again, the functional connectivity differences between the competition vs. cooperation condition and with the rTPJ seed, was correlating with each participant’s lying ratio in the competition conditions. The resulting significant regions, including rostral ACC (rACC), caudate, parahippocampal gyrus (PPA)/amygdala, and the IFG (see Figure 6, Table 5), were all negatively correlated. These results indicated that the lower the functional connectivities between the hub areas of different (e.g., mentalizing vs. reward-prediction) sub-networks, such as rTPJ-rACC, rTPJ-caudate, rTPJ-PPA/amygdala, etc; the higher probability the senders adopted the lying strategy. As additional checks, the dlpfc- and caudate-seed ROI PPI-deceiving rate correlation analyses both did not yield any significant clusters. These null findings further strengthen the singular importance of rTPJ seed in social deception behaviors.

**Figure 6.**
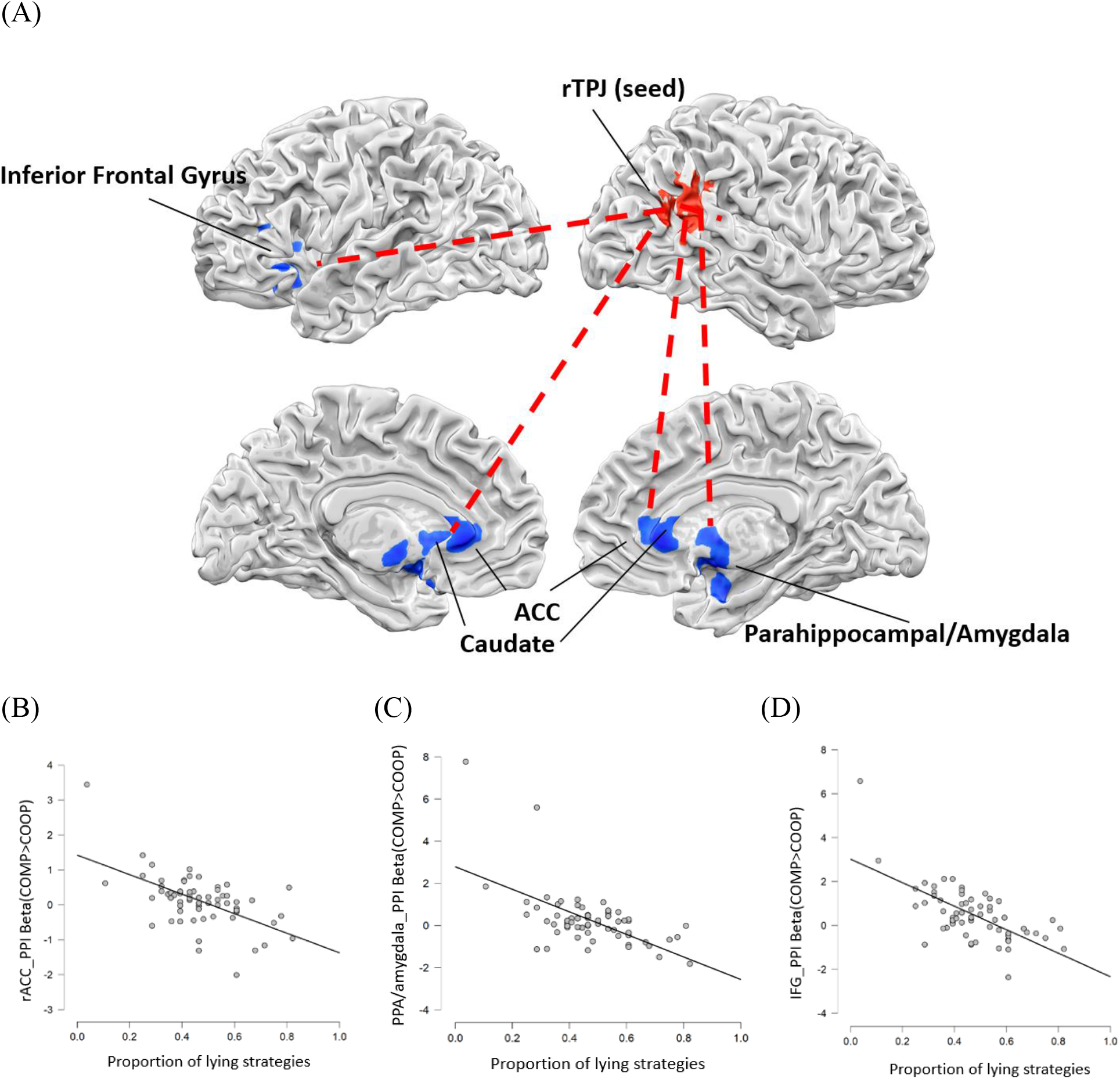
(A) The connectivity-based correlation map between rTPJ and other brain areas that showed highly negative correlations between their functional connectivity, and subjects’ proportion of lying strategies. (B)-(D): the scatterplots of individual’s lying ratio as senders and their connectivity measures; (B) rTPJ-rACC/caudate; (C) rTPJ-parahippocampal gyrus (PPA)/amygdala; and (D) rTPJ-IFG [Voxel-level threshold p < .005 uncorrected, (cluster threshold > 96 voxels, FWE p=.05 corrected thresholded), via Neuroelf’s Alphasim].

**Table 5.** With increase in participants’ proportion of lying strategies, the rTPJ (the seed) showed lower functional connectivity with the ACC, PPA/amygdala and IFG [Voxel-level threshold p < .005 uncorrected, (cluster threshold > 96 voxels, FWE p=.05 corrected thresholded), via Neuroelf’s Alphasim].

| Cluster | Anatomical region | x | y | z | correlation coefficient |
| --- | --- | --- | --- | --- | --- |
| Cluster1(Voxel : 120) | L Anterior Cingulate | 3 | 29 | 2 | -0.49 |
| Cluster2(Voxel : 111) | R Parahippocampal Gyrus (Amygdala) | 28 | -1 | -11 | -0.49 |
| Cluster3(Voxel : 100) | L Inferior Frontal Gyrus | -33 | 26 | -9 | -0.46 |

### Multi-Voxel Pattern Analysis (MVPA)

Given that our earlier GLM results showed no activation differences between truth-telling and lying strategies in the competition condition, it may be desirable to examine from another analysis viewpoint: whether the multi-voxel activation patterns of participants’ strategies under different conditions could be decoded by pattern classification schemes (Tsoi et al., 2016). Using the whole-brain searchlight mapping (Pereira & Botvinick, 2011), the group t-tests across 62 participants (minus two that never lied, and another two lied fewer than 5, out of the total 28, trials; and that each with p<.005 voxel level uncorrected, FWE p.05 cluster threshold 35 voxels) yielded ten ROIs that were commonly above-chance in classifying truth-telling vs. lying strategies in the competition condition. These 10 ROIs were: the rACC/vmPFC, angular gyrus, middle frontal gyrus (MFG), parahippocampal gyrus, lingual gyrus, IFG, middle temporal gyrus (MTG), putamen and the sub-gyral (see Fig.7A and Table 6 for details). Since each of the ROIs, despite being significant at the group level, exclusively reached only 55~56% classification accuracy, combining multiple (from one to ten ROIs, from the highest t-to the lowest t-values in the descending order) helped increase the classification accuracy to peak at 61.35%, hovering around the 5th-7th ROIs (Fig. 7B).

**Figure 7.**
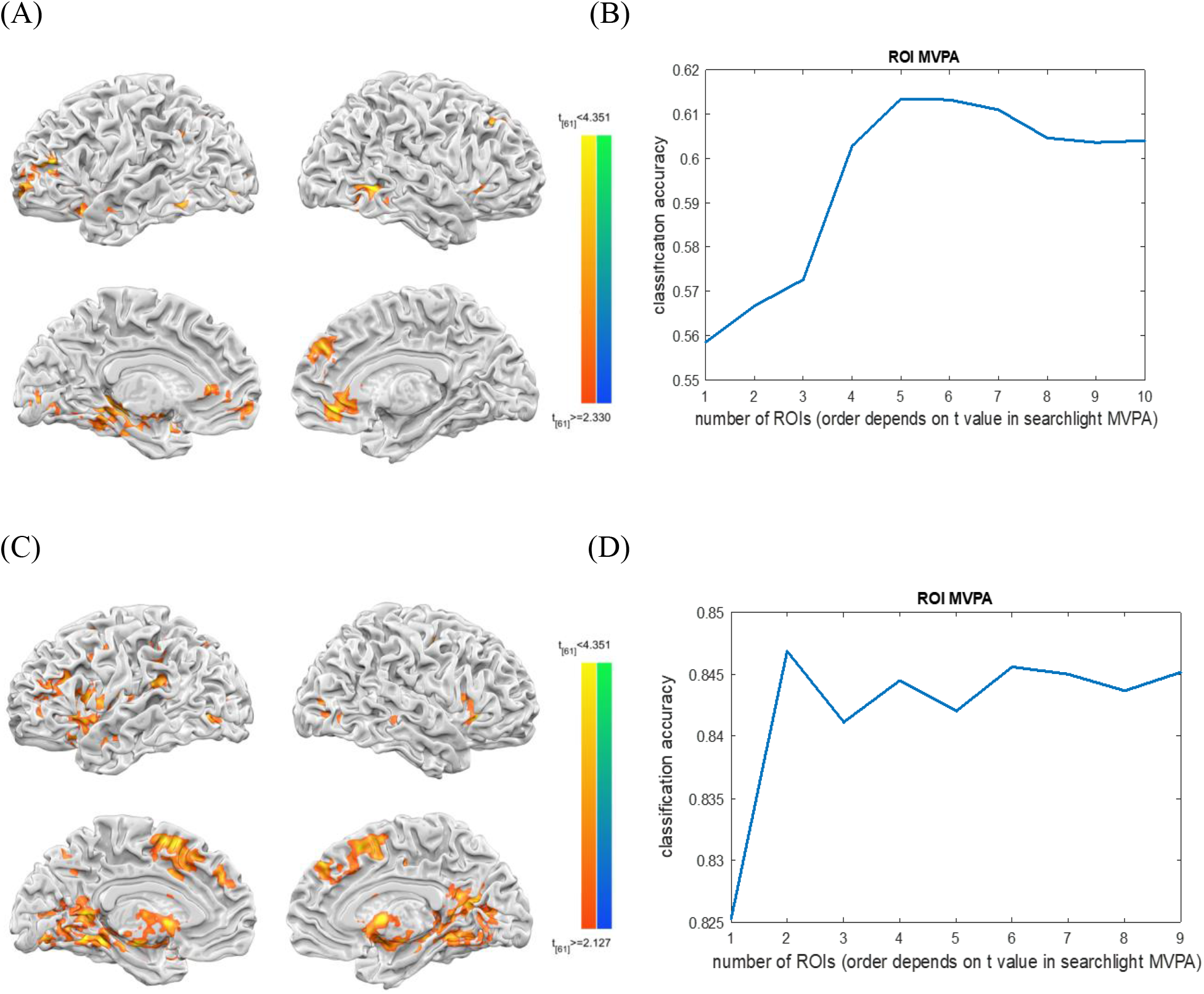
(A) Brain regions revealed for larger-than-chance performance (at the group level) in classifying truth-telling and lying strategies in the competition ($150) condition with the whole-brain multivariate searchlight mappings. The ten regions (t-values from top to bottom) included: the right rACC/vmPFC, the left angular gyrus, the left mPFC, the left parahippocampal gyrus, the left lingual gyrus, the left IFG, the right MTG, the right putamen and the left sub-gyral; (B) The average classification accuracy (N=62) in truth-telling vs. lying strategies under competition under accumulative ROI add-ups; (C) Brain regions revealed for >60% classification accuracy, at the group level, in classifying truth-telling in the cooperation ($200) versus in the competition ($150) conditions, via whole-brain searchlight MVPA. The nine ROIs included the SFG, lentiform nucleus, IFG, precentral gyrus, MFG, insula, IPL and precuneus; and (D) the average classification accuracy in truth-telling under the cooperation ($200) versus the competition ($150) conditions.

**Table 6.** Whole-brain searchlight MVPA in classification of truth-telling and lying strategies under competition [Voxel-level threshold p < .005 uncorrected, (cluster threshold >35 voxels, FWE p=.05 corrected thresholded), via Neuroelf’s Alphasim].

| Cluster | Anatomical region | x | y | z | t-value |
| --- | --- | --- | --- | --- | --- |
| Cluster1(Voxel : 100) | R Anterior Cingulate | 4 | 40 | -5 | 6 |
| Cluster2(Voxel : 79) | L Angular Gyrus | -39 | -61 | 34 | 5.54 |
| Cluster3(Voxel : 181) | L Middle Frontal Gyrus | -50 | 37 | 15 | 5.41 |
| Cluster4(Voxel : 514) | L Parahippocampal Gyrus | -37 | -43 | -6 | 5.38 |
| Cluster5(Voxel : 86) | L Lingual Gyrus | -16 | -81 | 0 | 5.31 |
| Cluster6(Voxel : 47) | L Inferior Frontal Gyrus | -26 | 20 | -14 | 4.72 |
| Cluster7(Voxel : 91) | R Middle Frontal Gyrus | 25 | 27 | 35 | 4.54 |
| Cluster8(Voxel : 73) | R Middle Temporal Gyrus | 48 | -58 | 5 | 4.46 |
| Cluster9(Voxel : 39) | L Sub-Gyral | -39 | -11 | -9 | 4.4 |
| Cluster10(Voxel : 50) | R Lentiform Nucleus<br>(Putamen) | 26 | 14 | 5 | 4.33 |

In contrast to the challenging classification results above, the same method was applied to the classification of truth-telling strategy in both the cooperation versus the competition condition. Because of its much stronger difference between two conditions (cf. Fig. 4B), the MVPA searchlight mappings have to shift the individual classification threshold, from the default 50% to 60%, to yield comparable clusters. The final group t-results listed nine ROIs, including the SFG, lentiform nucleus, IFG, precentral gyrus, MFG, insula, IPL and precuneus (see Fig. 7C and Table 7). When the first to the ninth ROIs were consecutively added for classification (‘truth-telling in cooperation vs. in competition condition’), the results peaked and hovered at 84.56 percent till the end (Fig. 7D).

**Table 7.** Whole-brain searchlight in classification of truth-telling under cooperation and competition [Voxel-level threshold p < .005 uncorrected, (cluster threshold >35 voxels, FWE p=.05 corrected thresholded), via Neuroelf’s Alphasim].

| Cluster | Anatomical region | x | y | z | t-value |
| --- | --- | --- | --- | --- | --- |
| Cluster1(Voxel : 490) | R Superior Frontal Gyrus | 11 | 19 | 50 | 5.64 |
| Cluster2(Voxel : 1835) | R Lentiform Nucleus | 24 | -20 | -1 | 5.28 |
| Cluster3(Voxel : 366) | L Inferior Frontal Gyrus | -42 | 17 | -10 | 4.84 |
| Cluster4(Voxel : 109) | R Precentral Gyrus | 35 | -4 | 43 | 4.72 |
| Cluster5(Voxel : 67) | R Middle Frontal Gyrus | 0 | 43 | 34 | 4.46 |
| Cluster6(Voxel : 115) | R Insula | 30 | 20 | 2 | 4.45 |
| Cluster7(Voxel : 130) | L Inferior Parietal Lobule | -44 | -41 | 40 | 4.28 |
| Cluster8(Voxel : 106) | L Precuneus | -19 | -62 | 36 | 4.21 |
| Cluster9(Voxel : 80) | R Middle Frontal Gyrus | -40 | 23 | 23 | 4.21 |

### Principal Component Analysis (PCA)

Given the null GLM contrast results in the truth-telling vs. lying in the competition ($150) condition, plus the merely ~61% group classification accuracy by adding multiple searchlight-identified ROIs (Fig. 7A and 7B), it may be tempting to adopt one or two dimension reduction algorithms to seek for opportunities to mitigate the noisy fMRI signals, thereby possibly improving this challenging classification. To this end, the principal component analysis, or PCA, widely considered a dimension reduction method to identify a few components that are good enough to classify both (a) truth-telling and lying strategies in competitions, and (b) truth-telling strategies in cooperation versus in competition conditions. The PCA analysis results suggested that when adding the top 150 components to classify truth-telling and lying strategies in the competition, the classification accuracy achieved 60% (Fig. 8A). In contrast, the classification accuracy could reach 80% by just combining the top 10 components to classify truth-telling strategies in the cooperation vs. in the competition conditions (Fig. 8B). To closely examine the individual variability behind Fig. 8A, Fig. 8C further to show individual participant’s classification performance (blue bars) under Fig. 7B and 8A (and the average performance rose to 74%, as the biased estimate), and the number of minimal principal components needed to achieve the best performance (red bars).

**Figure 8.**
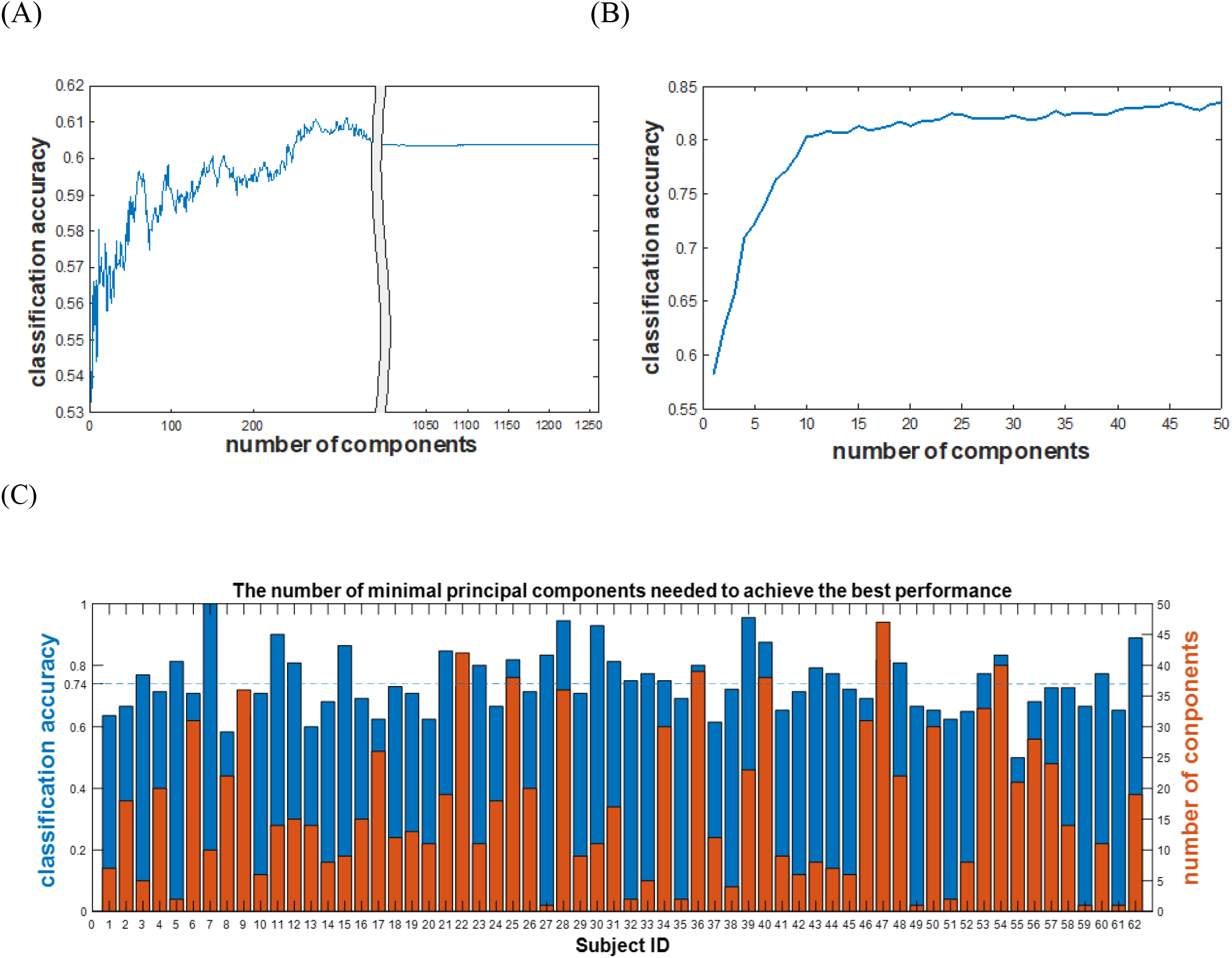
The result of principal component analysis (PCA): (A) when the ten searchlight-derived ROIs (totally 1260 voxels) were subjected to PCA, the ordered principal components were added up to predict truth-telling vs. lying strategies in the competition. The results suggested that when summed up to 150 components, classification accuracy reached 60% and peaked at 61% around 300 components; (B) when the nine searchlight-derived ROIs (totally 3298 voxels) entered the PCA, the resulting principal components also added up to predict truth-telling strategies in both the cooperation vs. competition conditions. The results showed that adding the first 10 components has already reached 80% accuracy, and the first 50 components, 84%; and (C) to highlight the individual variability in (A), here the best subject-level classification accuracy in predicting truth-telling and lying strategies in the competition condition for each participant (blue, and the average performance was 74%, as the biased estimate), and the corresponding minimal number of principal components needed (red), was shown.

## Discussion

The current study aims to investigate the neural underpinnings of social cooperation and deception with hyperscanning fMRI, a methodology that, despite dating back more than a decade ago (Montague et al., 2002), is still relatively scarce in the recent literature (Shaw et al., 2018; Spilakova, Shaw, Czekoova, & Brazdil, 2019; Xie et al., 2020). In the current study, a strategic opening-treasure-chest (OTC) game to solicit participants’ alternating responses as both senders and receivers, in which their strategies with the long-distance partner/opponents varied interactively. Under the equal expected value ($75) manipulation across two conditions, the 75% success rate in the cooperation trials (winners split the $200 reward) resulted in 98.2% cooperation truth-telling (sender) and 98.7% following (receiver) strategy in each pair. In contrast, the 50% success rate in the competition trials (winner took all the $150) rendered all participants adopting relatively equal rates of truth-telling vs. lying (as senders), and of following vs. un-following (as receivers). The behavioral responses mimicked the predictive pattern: almost 100% truth-telling and faithful following in the cooperation condition; 54.5% vs.45.5% in the ‘truth-telling vs. lying’ as senders, and 57.9% vs. 42.1% in ‘follow vs. unfollow’ as receivers of the competition condition; as well as longer response times in the competition, compared to those in the cooperation condition. In addition, the GLM results also align with the expected involvement of the 3 subnetworks: ToM network (e.g., rTPJ, mPFC, and precuneus) (Speer et al., 2020; Wang et al., 2020), reward processing (Abe & Greene, 2014), and executive control (Sun et al., 2017; Zhu et al., 2014), conforming to the hypotheses derived from the extant literature.

Indeed, the fact that our GLM contrasts showing three (ToM, executive/inhibition, and the reward processing) subnetworks simultaneously in a single study, may be the first when compared with similar fMRI/fNIRS studies that revealed idiosyncratic GLM mapping. For example, one study (e.g.,Yin & Weber, 2019), using the spontaneously lying strategy, incentivized by relatively high payoffs (honesty: $1, lying: $8, lying got caught: -$6) and low (probability = .2) caught-up rates, did not find the involvement of ToM network (see also Lisofsky et al., 2014, for meta-analysis results). Second, another mono-fMRI study (Volz et al., 2015; Zheltyakova, Kireev, Korotkov, & Medvedev, 2020), investigating lying in the social context through the inquiry of honest answers, and simple and sophisticated deceptions after each sender-receiver trial, did not find the executive/inhibition subnetwork in their GLM contrasts. Third, some recent fNIRS hyperscanning studies, through more naturalistic task designs (e.g., bluffing card game, and sender-receiver games) (Chen et al., 2020; Wang et al., 2020; Zhang et al., 2017), revealed both ToM and inhibition subnetworks, but was unable to probe into reward-related network (e.g., caudate nucleus), due to the depth limitation of NIRS (Ferrari & Quaresima, 2012; Pinti et al., 2020). Together, as Sip et al. (2008) reviewed, that although the three subnetworks were summarized, but most of the time selectively recruited in various forms of social deceptions, our GLM findings, when compared with the extant literature, additionally suggest the importance of both task designs and research tools that contribute to the patterns of analysis results.

The joint mappings of precuneus region when overlaying of functional connectivity results from the rTPJ-, the DLPFC-, and the caudate-seed, all comparing competition vs. cooperation conditions (Fig. 5), given its rarity in the literature, strongly suggest a compelling point: the 3 subnetworks (execution/inhibition: DLPFC; reward processing: caudate, and ToM: rTPJ) are not commonly recruited (as shown in GLM contrast, Fig. 4A) together, but are also functionally, even simultaneously, engaged, during the task trials. The commonly connected region precuneus has been implicated, both structurally and functionally (Cunningham et al., 2017), as one of the vital hub regions (van den Heuvel & Sporns, 2013) of the default mode network (Utevsky, Smith, & Huettel, 2014). These lines of evidence, combined with the similar PPI results in terms of rTPJ-seed connectivity findings, emphasize once again the intricate interactions/integrations of ToM, reward, and execution subnetworks, almost in every trial of competition condition, regardless of truth telling or lying trials.

Other than the common findings between GLM contrasts and PPI results of the similar conditions (Fig. 4 and 5), the individual (negative) correlations between the behavioral lying rates and their functional connectivity between the rTPJ (seed region) and rACC/caudate, parahippocampal gyrus/amygdala and IFG (shown in Fig. 6), also provide novel perspectives into these results. These connectivity-based correlations, along with the null findings with other DLPFC-nor caudate-seeds, suggest the common involvement of these subnetworks (e.g., rTPJ-IFG belong to ToM and inhibition networks separately; and rTPJ-hippocampus and -amygdala are for ToM and event memories, episodic emotions, etc) in predicting individual strategic tendencies. Similar with the practices in Hackel et al. (2020) and Speer et al. (2020), here we term this procedure ‘Individual Connectivity Correlation’, or ICC, different from the common practices of whole-brain correlations between condition-wise beta estimates and behavioral covariates (Abe, 2011; Suzuki et al., 2015; Wu et al., 2011). While the latter practices will likely continue, a recent preprint has suggested the reasonable sample size of brain-wide association studies to be 2000 (Marek et al., 2020), a number that seems both daunting and implicative. In light of recent suggestions of the connectivity-related measures more reliable (than the ROI-based ones) (Elliott et al., 2020), the prospect of these ICCs being more reliable/reproducible remains to be verified.

The ROI MVPA analysis, in light of the earlier success of revealing subtle (but significant) pattern differences on the null findings by univariate GLM (Tsoi et al., 2016), was subsequently deployed by combining a growing number (from ROI_1 to ROI_10, with descending group wise t-values) of searchlight-based ROIs of similar contrasts (e.g., Fig. 7A for truth-telling vs. lying in the competition condition). As Fig. 7B showed, the multiple-ROI MVPA could only raise the accuracies to around 61% with 5~6 ROIs (shown in Fig. 7A, with decreasing t-values). Although reaching what was expected (seeking significant pattern difference out of null univariate findings), these results were in sharp contrast to the later comparisons of truth-telling in either cooperation or in competition conditions (Fig. 7C and 7D, 84.5% with two ROIs, and fluctuated only mildly). These results can be restated as “telling truth-telling in different scenarios, through the manipulation of different incentive rules, is relatively easy; but telling whether someone is truth-telling or lying in the same competition context, wherein the same incentive rules applies, is hard”. The latter condition, classifying truth-telling from lying in the competition condition, is also related to the sophisticated deception (Volz et al., 2015), second-order deception (Ding, Sai, Fu, Liu, & Lee, 2014), or the famous ‘Empty Fort Strategy’, 32th of the 36 Chinese Stratagems. Despite inquiring participants about their underlying motivation after each answer, Volz et al. (2015) still faced the risk of participants not being totally honest about their responses. Similarly incapable in the current study, we did however create two experimental contrasts where the same name ‘deception’ could be contextually differentiated, based on the interactions among reward probabilistic, reward dissemination rules, individual characteristics, etc.

Lastly, as a general fix to the overfitting and multicollinearity problems in multivariate analyses, principal component analysis (PCA) has been, and was adopted in the current study, to reduce the number of dimensions from high-dimensional MVPA features/voxels. The results demonstrated that, when collapsing all the nine ROIs (see Fig. 7C) which consistently showed larger than 80% classification of “truth-telling in cooperation vs. that in competition” (Fig. 7D), after PCA each ROI can be reduced to <10 components (Fig. 8B). However, classifying truth-telling vs. lying strategies in competition required ~150 components (Fig. 8A) to achieve the same (~60%) classification performance (Fig. 7B). The further examination in Fig. 8C, showing different numbers of PCA components (red, right) to achieve the best classification performance (blue, left) in predicting truth-telling and lying strategies. Therefore, “classifying truth-telling in the cooperation vs. in the competition” versus “classifying truth-telling vs. lying in the competition condition” is not only ‘easy (84.5) vs. hard (61%)’, but also different in terms of ‘number of optimal components’ (~10 vs. ~150), an indirect indicator of increasing individual variability, which is reflected in Fig. 8C. The boosted 74% in Fig. 8C, compared to the original 61% (c.f. Fig. 7B), though a biased maximization, nonetheless manifested participant idiosyncrasy from another perspective.

## Limitations

One of the limitations is the incapability of the current design to truly separate the mental strategy behind truth-telling vs. lying behaviors in the competition condition. Future versions of the sender-receiver Open-Treasure-Chest game could adopt a condition of 100% reward (on which side) condition, similar to the dyad bluffing game (Zhang et al., 2017), to better separate the true truth-telling from sophisticated deception (Volz et al., 2015), from the same truth-telling behavior under the competition condition. In addition, whether the within-subject behavioral tendencies tend to go hand in hand, (e.g. are trustworthy senders also faithful followers?) And how that affects the data analysis scheme (Wang et al., 2020) and result interpretation, are behind the scope of this study, and awaiting future endeavors.

## Conclusion

Four (univariate, seed-based connectivity, multivariate, and dimension-reduction) data analysis approaches were applied to a two-person hyperscanning fMRI experiment. Three types of neural networks: ToM, inhibition, and reward processing, were iteratively involved in both the cooperation and competitions, and some of the inter-connectivity tied to their strategic tendencies. Conditions of interest greatly affect the classification accuracy. We argue that instead of grace/will hypotheses (Greene & Paxton, 2010) by default, people are more influenced by contextual manipulations (e.g., different awarding rules for $200 vs. $150), and integrate the relative benefit of self vs. other, event histories, and the outcome probabilities, to reach optimal decisions.

## Acknowledgments

This work is supported by MOST 107-2420-H-006-007-MY3. We thank Mind Research and Imaging Center (MRIC) at National Cheng Kung University for consultation and instrument availability.

